# Fentanyl depression of respiration: comparison with heroin and morphine

**DOI:** 10.1101/662627

**Authors:** Rob Hill, Rakulan Santhakumar, William Dewey, Eamonn Kelly, Graeme Henderson

## Abstract

**Background and Purpose:** Fentanyl overdose deaths have reached ‘epidemic’ levels in North America. Death in opioid overdose invariably results from respiratory depression. In the present work we have characterized how fentanyl depresses respiration and by comparing fentanyl with heroin and morphine, the active breakdown product of heroin, we have sought to determine whether there are factors, in addition to high potency, that contribute to the lethality of fentanyl.

**Experimental Approach:** Respiration (rate and tidal volume) was measured in awake, freely moving mice by whole body plethysmography

**Key Results:** Intravenously administered fentanyl produced more rapid depression of respiration than equipotent doses of heroin or morphine. Fentanyl depressed both respiratory rate and tidal volume, the effect on tidal volume may reflect increased respiratory muscle stiffness. Fentanyl did not depress respiration in μ opioid receptor knock-out mice. Naloxone, the opioid antagonist widely used to treat opioid overdose, reversed the depression of respiration by morphine more readily than that by fentanyl whereas diprenorphine, a more lipophilic antagonist, was equipotent in reversing fentanyl and morphine depression of respiration. Prolonged treatment with morphine induced tolerance to respiratory depression but the degree of cross tolerance to fentanyl was less than the tolerance to morphine itself.

**Conclusion and Implications:** We propose that several factors (potency, rate of onset, muscle stiffness, lowered sensitivity to naloxone and lowered cross tolerance to morphine) combine to make fentanyl more likely to cause opioid overdose deaths than other commonly abused opioids.

## INTRODUCTION

Since 2013 there has been a dramatic rise in acute opioid overdose deaths involving new synthetic opioids, primarily the fentanyls (fentanyl and structurally related medicinal and illicit drugs), in North America (NIH, 2019). Of the over 60,000 opioid overdose deaths in the USA in 2017 almost 30,000 involved fentanyls, exceeding those involving heroin or prescription opioids such as oxycodone and hydrocodone. Elsewhere, in Europe, fentanyl deaths have been recorded in Estonia (for some time fentanyls were the main street opioids available in that country), and there have been sporadic outbreaks of fentanyl-related deaths in the UK, Germany and Finland (EMCDDA, 2018). Ease of synthesis (cf. the need to grow swathes of opium poppies to produce heroin), high potency (smaller quantities need to be shipped by comparison with heroin) and ease of purchase on the dark web make the fentanyls attractive to suppliers of illicit opioids. Fentanyls are frequently mixed with heroin to increase its potency (Griswold *et al.*, 2017; Marinetti *et al.*, 2014). A recent development is the addition of fentanyls to cocaine products and to illicit prescription opioid and benzodiazepine tablets (Green *et al.*, 2016; Sutter *et al.*, 2017). Death in fentanyl overdose results primarily from depression of respiration (Mathers *et al.*, 2013; Pierce *et al.*, 2015). Fentanyls and other opioid agonists depress respiration by acting on μ opioid receptors at various sites to reduce the response to raised pCO_2_ and lowered pO_2_ and thus reduce the drive to breathe (Pattinson, 2008). This reduction in respiratory drive results in a decrease in the rate of breathing and in periods of apnea (cessation of breathing) which in extremis results in death.

A number of factors may contribute to why the fentanyls are so deadly. Their high potency means that only small amounts are required to produce profound effects and thus even a small error in weighing out the drug can result in too much being taken. Rapid penetration into the brain can result in overdose levels being reached more quickly than with heroin. Deaths in heroin overdose may take more than 30 min to occur after injection (Darke *et al.*, 2016), providing a window of opportunity for intervention (administration of the opioid antagonist naloxone). In contrast, fentanyl overdose deaths can occur very quickly before there is an opportunity to administer naloxone (Burns *et al.*, 2016). Fentanyls induce muscle stiffness (Benthuysen *et al.*, 1986; Streisand *et al.*, 1993) including in intercostal and diaphragm muscles, often referred to as ‘wooden chest’, and this is likely to make it harder to breathe. There have been several reports suggesting that depression of respiration by fentanyls shows reduced sensitivity to reversal by naloxone (Fairbairn *et al.*, 2017; Lynn *et al.*, 2018; Peterson *et al.*, 2016; Schumann *et al.*, 2008). In one report, several cases were recorded in which multiple doses of naloxone were required before recovery of respiration following a fentanyl overdose (Somerville *et al.*, 2017). Furthermore, a low level of cross tolerance between heroin and fentanyls may allow the fentanyls to retain potency to depress respiration in individuals tolerant to heroin-induced respiratory depression. In the present paper we have characterized how fentanyl depresses respiration in mice and have sought to determine how the factors described above may contribute to fentanyl overdose.

## METHODS AND MATERIALS

### Animals

Male CD-1 mice weighing approximately 30 g were obtained from Charles River (UK). Breeding pairs of μ opioid receptor knock-out mice (*Oprm1*^*tm1Kff*^) were obtained from the Jackson Laboratory (USA) and offspring bred in house at the University of Bristol. Both male and female μ opioid receptor knock-out mice weighing between 25 and 30 g were used in this study. Male and female wild type C57BL/J mice weighing approximately 30 g were purchased from Charles River (UK). All mice were maintained at 22 °C on a reversed 12 h dark-light cycle with food and water available *ad libitum*. All experiments were performed in the dark (active) phase. Mice were randomly ascribed to treatment groups with the experimenter blinded to the drug treatment or knock-out/wild type phenotype as appropriate.

All procedures were performed in accordance with the UK Animals (Scientific Procedures) Act 1986, the European Communities Council Directive (2010/63/EU) and the University of Bristol ethical review document, as well as the ARRIVE and British Journal of Pharmacology guidelines.

### Measurement of respiration

Respiration was measured in freely moving mice using plethysmography chambers (EMKA Technologies, France) supplied with a 5% CO_2_ in air mixture (BOC Gas Supplies, UK) as described previously (Hill *et al.*, 2016). Rate and depth of respiration were recorded and averaged over 1 or 5 min periods (except immediately after drug injection when the time period was 0.5 or 3 min respectively) and converted to minute volume (rate x tidal volume). Drugs were administered i.v. or i.p. as indicated.

### Measurement of locomotor activity

A beam break rig (Linton Instrumentation, UK) was used to assess the locomotor activity of mice. An automated data logging suite (AMON Lite, Linton Instrumentation, UK) was used to track the movement of mice throughout the experimental session. On the day prior to locomotor assessment each mouse was placed in a fresh cage and allowed to explore the cage for 30 min. On the experimental day the mouse was again allowed to explore the cage for 30 min before drug administration. Locomotor activity was then measured for 30 min following drug administration. Mice had access to water *ad libitum* but had no access to food in either session in order to dissuade rearing and climbing behaviour.

### Drug administration

In most experiments we have used i.p. injection as this is a simple and relatively non-stressful means of drug administration to mice. However, in those experiments designed to determine rate of onset of effect opioid agonist drugs were administered i.v. with mice restrained in a clear plastic tube while opioid or vehicle was administered by tail vein injection in a 0.1 ml volume.

### Reversal of opioid respiratory depression by opioid antagonists

Respiration was recorded for 20 min followed by an acute i.p. injection of opioid or vehicle. Respiration was then recorded for 20 min following opioid/vehicle administration, allowing maximal depression of respiration to occur. Naloxone or diprenorphine was then administered (20 min after opioid/vehicle) by i.p. injection on the opposite side of the peritoneal cavity to the opioid/vehicle injection.

### Induction of morphine tolerance

Tolerance to morphine respiratory depression was induced by 3x 100 mg.kg^−1^ morphine injections, administered 12 h apart followed by subcutaneous implantation of morphine-filled osmotic mini-pumps delivering 45 mg.kg^−1^.day^−1^ morphine for a total of 6 d as described previously (Hill *et al.*, 2018; Hill *et al.*, 2016). Control mice were injected with saline and implanted with minipumps that were filled with saline.

### Experimental design and data analysis

Data from previous experiments where respiratory depression and locomotor activity were measured either following acute opioid administration in naïve mice or following pump implantation were subjected to post hoc power analyses using G*Power (version 3.1.9). Our calculations indicated that for depression of respiration n=6 (acute experiments) or n=7 (pump experiments) and for locomotor activity n=8 for each individual group would produce a significant result if an actual effect occurred.

For each mouse the change in each behavioural parameter (respiratory rate, tidal volume, minute volume and locomotor activity) following acute drug administration has been calculated as the percentage of the pre-drug baseline as described previously (Hill *et al.*, 2018; Hill *et al.*, 2016; Withey *et al.*, 2017). Area under the response versus time curve (AUC) was determined using a 100% baseline as described previously (Hill *et al.*, 2016). Overall changes from a single factor were analysed using a One-way ANOVA with Bonferroni’s post-test. GraphPad Prism 5 was used for all statistical analyses. All data are displayed as mean ± standard error of the mean (s.e.m.). The data and statistical analyses comply with the recommendations on experimental design and analysis in pharmacology (Curtis *et al.*, 2015).

### Drugs and chemicals

Heroin hydrochloride (diacetyl morphine hydrochloride), morphine hydrochloride (both from Macfarlane Smith, UK), fentanyl citrate, naloxone hydrochloride, naltrindole hydrochloride and norbinaltorphimine dihydrochloride (all from Sigma Aldrich, UK) were dissolved in sterile saline. Diprenorphine (Tocris, UK) was dissolved in water by adding an equivalent amount of hydrochloric acid and subsequently diluted in sterile saline.

### Nomenclature of targets and ligands

Key protein targets and ligands in this article are hyperlinked to corresponding entries in http://www.guidetopharmacology.org, the common portal for data from the IUPHAR/BPS Guide to PHARMACOLOGY (Harding *et al.*, 2018), and are permanently archived in the Concise Guide to PHARMACOLOGY 2017/18 (Alexander *et al.*, 2017).

## RESULTS

### Depression of respiration by fentanyl, heroin and morphine

To study the rate of onset of respiratory depression fentanyl, heroin and morphine, were administered i.v. to mice. Morphine (7.5 mg.kg^−1^), heroin (7.5 mg.kg^−1^) and fentanyl (112 μg.kg^−1^) each depressed minute volume significantly, reaching approximately the same degree of respiratory depression 10 - 15 min after injection (Fig. 1A). Saline injection did not significantly alter respiration. The half-time to reach maximum depression of minute volume was determined for each agonist by fitting a single exponential curve (Fig. 1A). Fentanyl had the fastest rate of onset and morphine the slowest (Table 1). The rate of onset of respiratory depression correlated with the lipophilicity of each drug (Table 1).

**Table 1.** Rate of onset of opioid depression of respiration following i.v. administration and calculated lipid solubility values (ClogP) for opioid agonists and antagonists. The data for the onset of respiratory depression for opioid agonists were fitted to a single exponential (see Fig. 1A) to obtain the t_1/2_ value for each opioid drug. Data are presented as the mean ± s.e.m; n = 12 for each drug. Statistical comparison was made by 1-way ANOVA with Bonferroni’s comparison. * indicates statistical difference (p<0.05) from morphine and ¶ indicates statistical difference (p<0.05) from heroin [F = 5.093 (dfn = 2, dfd = 36)]. The ClogP value for each drug was calculated using Chem3D (PerkinElmer).

| Opioid | Rate of onset<br>($t_{1/2}$, min) | ClogP |
| --- | --- | --- |
| Fentanyl | $0.54 \pm 0.09^{*\P}$ | 3.62 |
| Heroin | $1.70 \pm 0.68^*$ | 1.48 |
| Morphine | $4.64 \pm 1.62$ | 0.57 |
| Naloxone | - | 0.16 |
| Diprenorphine | - | 2.66 |

**Figure 1.**
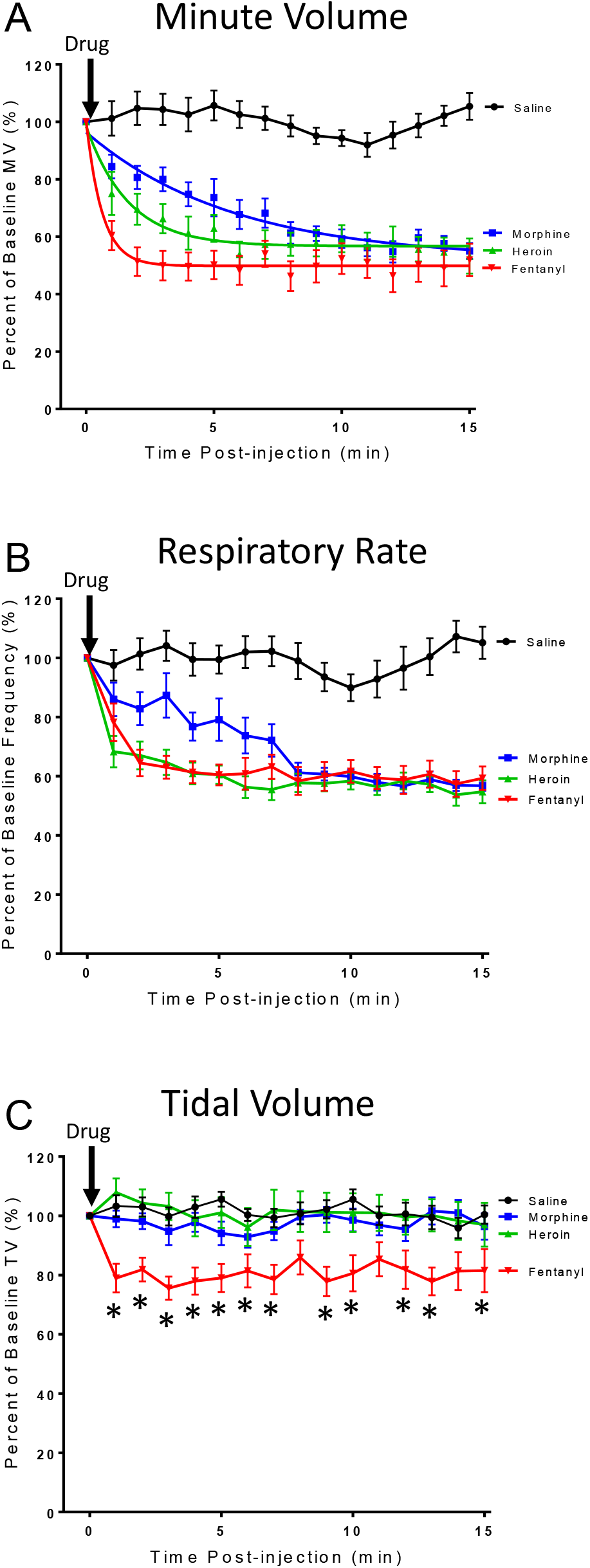
Rate of onset of opioid respiratory depression. Respiratory parameters were monitored in mice receiving i.v. injection of fentanyl (112 μg.kg^−1^), heroin (7.5 mg.kg^−1^) morphine (7.5 mg.kg^−1^) or saline. **A.** Fentanyl, heroin and morphine rapidly depressed minute volume (MV), the effect of the drugs reaching a similar steady state 10 - 15 min post-administration. Data for each drug are fitted to a single exponential. **B.** Fentanyl, heroin and morphine depressed respiratory rate. **C.** Heroin and morphine had no effect on tidal volume (TV), whereas fentanyl significantly depressed tidal volume [F = 65.05 (dfn = 3, dfd = 704)]. **A-C.** Saline injection did not alter any of the respiratory parameters. All data presented as mean ± s.e.m. Statistical comparison in **C** was made by 2-way ANOVA with Bonferroni’s comparison. * indicates p<0.05 compared to saline. n=12 for each group.

Both morphine and heroin depressed respiration by decreasing the rate of respiration, having no effect on tidal volume (Fig. 1B & C). As previously reported (Hill *et al.*, 2018; Hill *et al.*, 2016), tidal volume was maintained due to the prolongation of inspiration by apneustic compensation. However, fentanyl depression of respiration resulted from both a decrease in the rate of respiration (Fig. 1B) and a decrease in tidal volume (Fig. 1C). As the rate of onset of the decrease in rate of respiration was the same for both heroin and fentanyl (Fig. 1B) then the faster depression of minute volume by fentanyl (see Fig. 1A) seems likely to be due to the rapid decrease in tidal volume (Fig. 1C).

We have previously demonstrated the dose-dependency of depression of respiration by morphine and oxycodone in mice breathing 5% CO_2_ in air (Hill *et al.*, 2018; Hill *et al.*, 2016). Heroin (1 - 90 mg.kg^−1^ i.p.) produced a dose-dependent depression of minute volume and respiratory rate (Fig. 2A, E & G) but only slightly decreased tidal volume at the highest dose of 90 mg.kg^−1^ (Fig. 2C, F & G). The potency of heroin to depress minute volume was similar to that previously reported for morphine (Hill *et al.*, 2016). Fentanyl (0.05 - 1.35 mg.kg^−1^ i.p.) produced a dose-dependent depression of tidal volume, respiratory rate and minute volume (Fig. 2B, D, E, F & G). To depress minute volume fentanyl was approximately 70-fold more potent than heroin.

**Figure 2.**
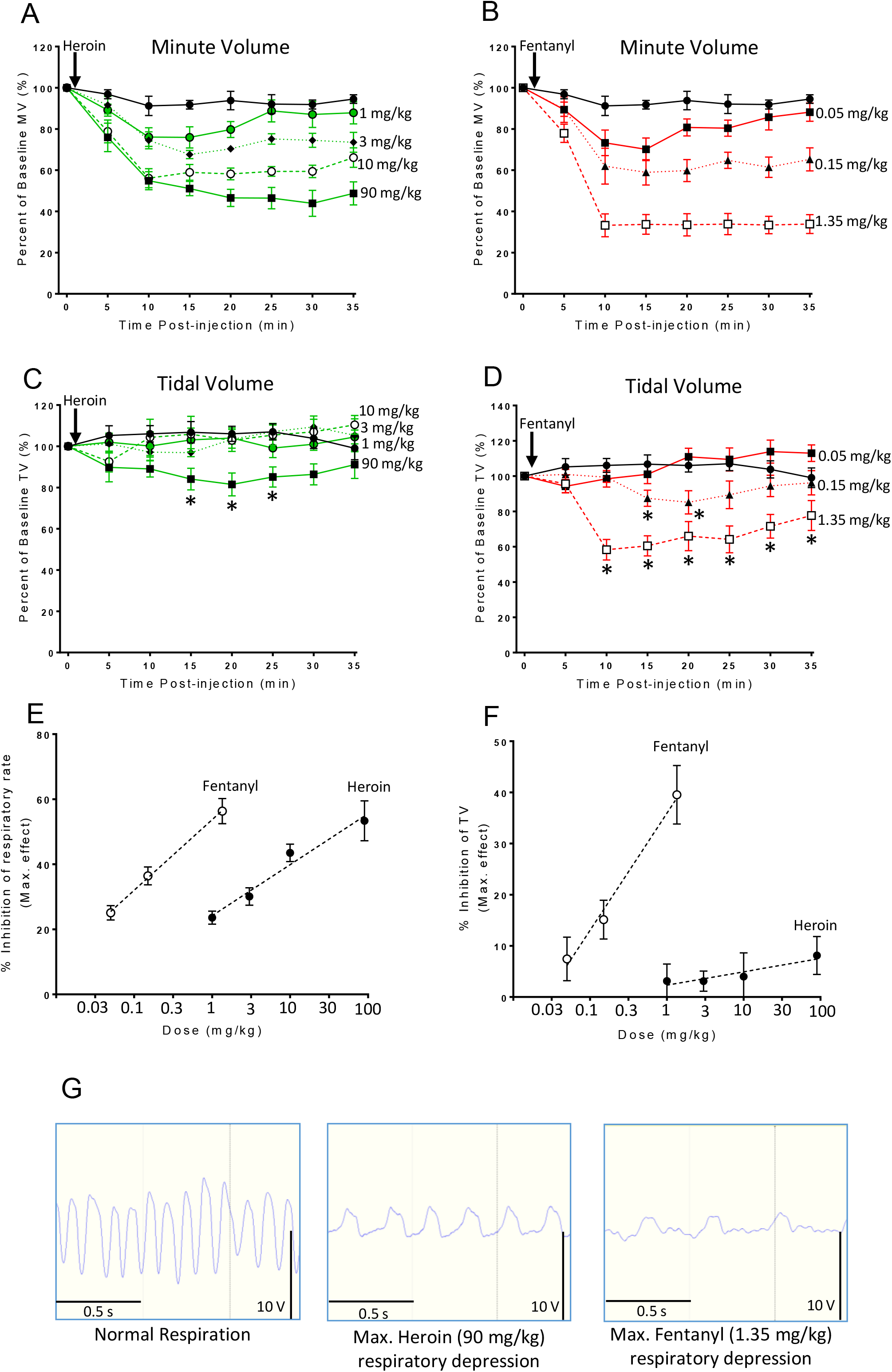
Effect of heroin or fentanyl on mouse respiration. **A.** Heroin (1 - 90 mg.kg^−1^, i.p.) dose-dependently depressed minute volume (MV). **B.** Fentanyl (0.05 - 1.35 mg.kg^−1^, i.p.) dose-dependently depressed minute volume. **C.** Heroin only slightly depressed tidal volume (TV) at the highest dose tested. **D.** Fentanyl dose-dependently depressed tidal volume. In **C** [F = 13.07 (dfn = 4, dfd = 192)] & **D** [F = 58.33 (dfn = 3, dfd = 160)] statistical comparison was made by 2-way ANOVA with Bonferroni’s comparison. * indicates p<0.05 compared to saline pre-treated mice. n=6 for each group. In **A** & **B** statistical significance was observed for most time points for each dose of heroin and fentanyl but *s have been omitted for clarity as the size of effect of each dose is clear from the dose-response graphs in **E** & **F**. **E**. Dose-response curves for fentanyl and heroin depression of respiratory rate (data calculated from experiments shown in A & B). **F**. Dose-response curves for fentanyl and heroin inhibition of tidal volume (data calculated from experiments shown in C & D). **G.** Left hand trace - control respiratory trace in the absence of opioid. Middle trace – at maximum respiratory depression by heroin 90 mg.kg^−1^. Right hand trace - at maximum respiratory depression by fentanyl 1.35 mg.kg^−1^.

### Effects of fentanyl and heroin on locomotor activity

In mice, at high doses, morphine and other μ opioid receptor agonists increase locomotor activity (Hill *et al.*, 2016; Lessov *et al.*, 2003; Valjent *et al.*, 2010). In the present study we observed that heroin (90 mg.kg^−1^ i.p.) increased locomotor activity but in contrast, fentanyl (1.35 mg.kg^−1^ i.p.) decreased locomotor activity (Fig. 3).

**Figure 3.**
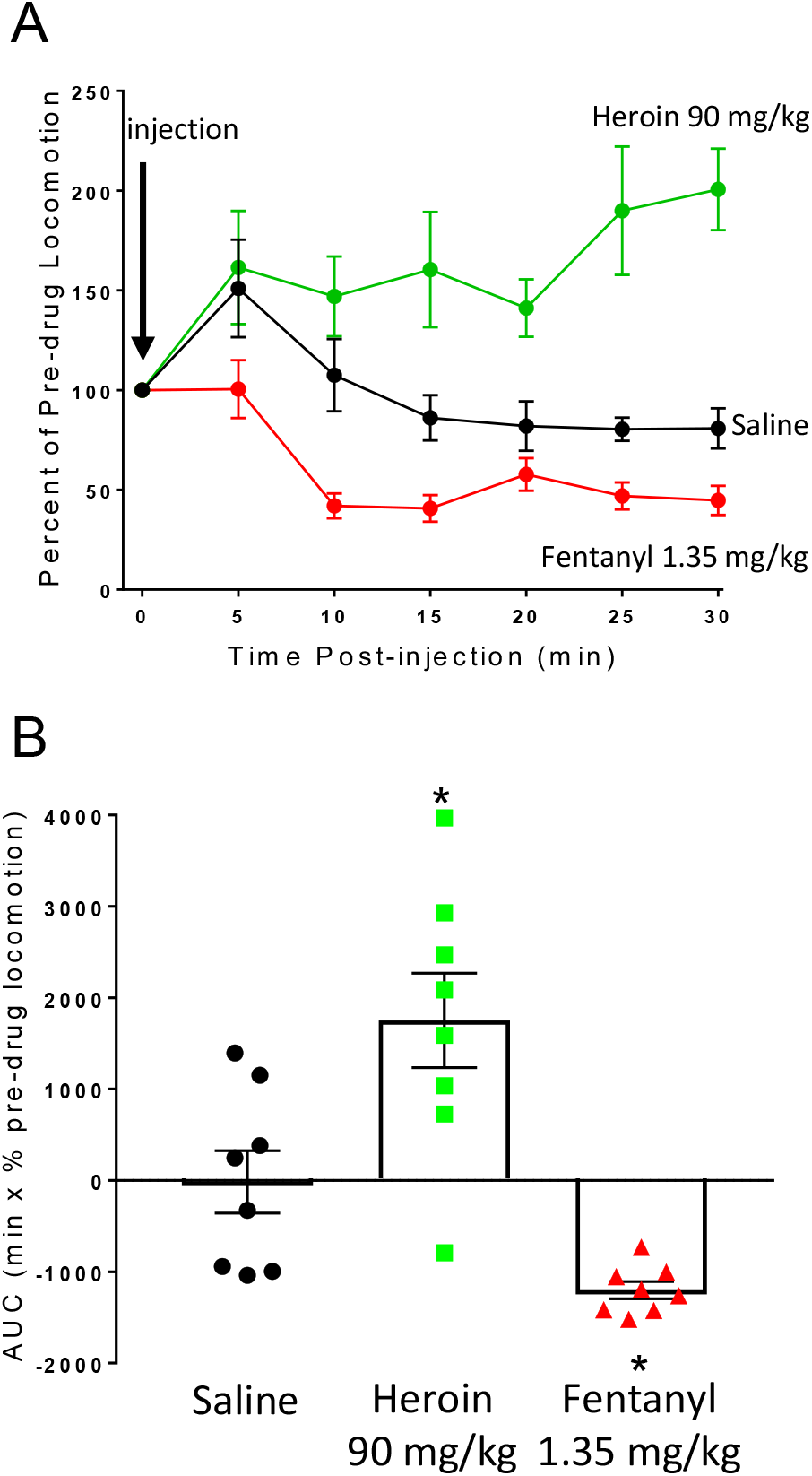
Change in mouse locomotor activity following heroin or fentanyl administration. **A.** Saline (i.p.), heroin (90 mg.kg^−1^ i.p.) or fentanyl (1.35 mg.kg^−1^ i.p.) were administered to mice and locomotor activity measured. Heroin caused a sustained increase in locomotor activity compared to saline, whereas fentanyl caused a decrease in locomotor activity compared to saline. **B.** Area under the curve (AUC) analysis of data in **A.** Statistical comparison in **B** made using 1-way ANOVA with Bonferroni’s comparison. * indicates p<0.05 compared to saline. n=8 for each group.

### Antagonism of fentanyl respiratory depression

It has been suggested that in overdose in humans the depression of respiration by fentanyl is less effectively reversed by naloxone compared to that by heroin (Peterson *et al.*, 2016; Schumann *et al.*, 2008; Somerville *et al.*, 2017). To investigate antagonist reversal of opioid respiratory depression, we administered equipotent doses of morphine, the main active breakdown product of heroin, and fentanyl (10 mg.kg^−1^ and 0.15 mg.kg^−1^ i.p. respectively) to mice, allowed maximal depression of respiration to develop over 20 min and then administered naloxone or diprenorphine. Naloxone (0.3 mg.kg^−1^ i.p.) rapidly antagonised the depression of respiration induced by morphine, with full reversal being apparent 5 min after naloxone administration (Fig. 4A). In contrast, the same dose of naloxone did not produce only slight reversal of the respiratory depression induced by fentanyl (Fig. 4A). Naloxone (1 mg.kg^−1^ i.p.) also fully reversed respiratory depression induced by morphine, but only partially reversed that induced by fentanyl (Fig. 4B). Only when naloxone (3 mg.kg^−1^ i.p.) was administered did full reversal of the respiratory depression induced by fentanyl occur (Fig. 4C), i.e. a 10-fold greater dose of naloxone was required to reverse fentanyl respiratory depression compared to that by an equi-potent dose of morphine. In contrast to naloxone, diprenorphine (0.03 mg.kg^−1^ i.p.) partially reversed equipotent doses of morphine and fentanyl to the same degree, though some respiratory depression persisted with both agonists (Fig. 4D). However, administration of diprenorphine (0.09 mg.kg^−1^ i.p.) rapidly and fully reversed both morphine and fentanyl depression of respiration (Fig. 4E).

**Figure 4.**
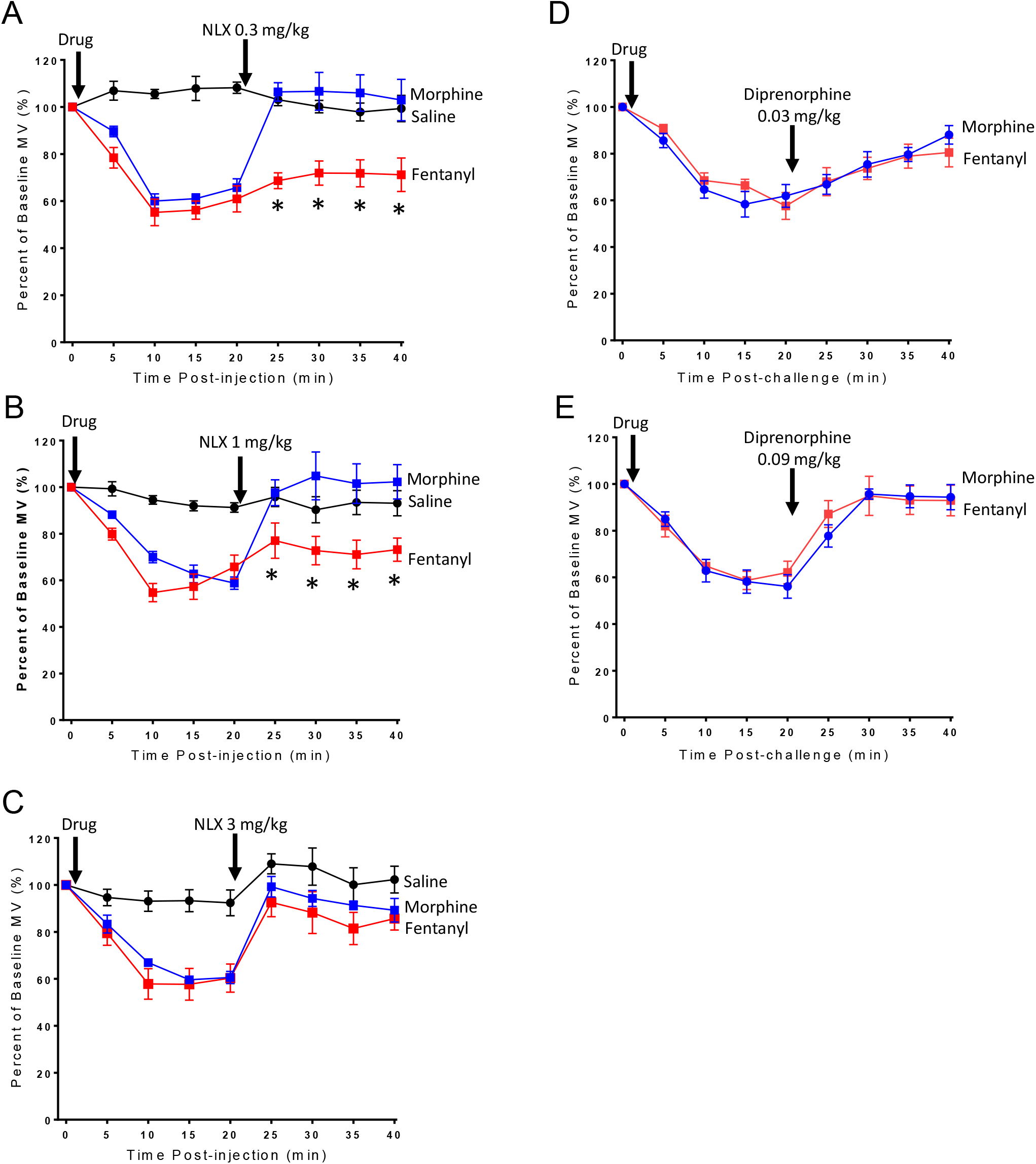
Reversal of morphine and fentanyl respiratory depression by naloxone and diprenorphine. **A-C** Equipotent respiratory depressant doses of morphine (10 mg.kg^−1^ i.p.) and fentanyl (0.15 mg.kg^−1^ i.p.) were administered to mice before naloxone administration 20 min after saline or opioid agonist. **A.** Naloxone 0.3 mg.kg^−1^ i.p. fully reversed morphine respiratory depression, but fentanyl respiratory depression was unaffected [F = 55.3 (dfn = 2, dfd = 15)]. **B.** Naloxone 1 mg.kg^−1^ i.p. fully reversed morphine respiratory depression, whereas fentanyl was only partially reversed [F = 14.98 (dfn = 2, dfd = 15)]. **C.** Naloxone 3 mg.kg^−1^ i.p. fully reversed both morphine and fentanyl respiratory depression. **D.** Diprenorphine 0.03 mg.kg^−1^ i.p. partially reversed morphine and fentanyl respiratory depression to the same degree **E.** Diprenorphine 0.09 mg.kg^−1^ i.p. fully and equally reversed both morphine and fentanyl respiratory depression back to baseline levels. All data presented as mean ± s.e.m. Statistical comparison of minute volume following naloxone administration made by 2-way ANOVA with Bonferroni’s comparison. * indicates p<0.05 compared to saline. n=6 for each group.

Fentanyl has been reported to be a relatively selective agonist at μ opioid receptors showing 100-fold and 400-fold higher affinity for the μ opioid receptor over κ and δ opioid receptors respectively (Toll *et al.*, 1998). To examine the possibility that there may be a δ opioid receptor component to fentanyl’s respiratory depressant activity we have assessed the ability of the δ opioid antagonist naltrindole to prevent the effect of fentanyl on minute volume and tidal volume in CD-1 mice. Pre-treatment with naltrindole (10 mg.kg^−1^) failed to prevent the depression of either minute volume or tidal volume produced by fentanyl (1.35 mg.kg^−1^); indeed, naltrindole slightly potentiated the effect of fentanyl (Fig. 5A & B) which suggests that endogenous opioid tone through δ opioid receptors can modulate the action of fentanyl. Furthermore, pre-treatment of CD-1 mice with the κ opioid receptor antagonist NorBNI (10 mg.kg^−1^ i.p.) 24 h prior to fentanyl administration did not affect the respiratory depressant effects of fentanyl (1.35 mg.kg^−1^) (Fig. 5C & D). We have previously reported that this treatment with NorBNI prevented the antinociceptive response to the κ opioid receptor agonist U69593 in the tail flick latency assay and that U69593 alone did not depress respiration (Hill *et al.*, 2018).

**Figure 5.**
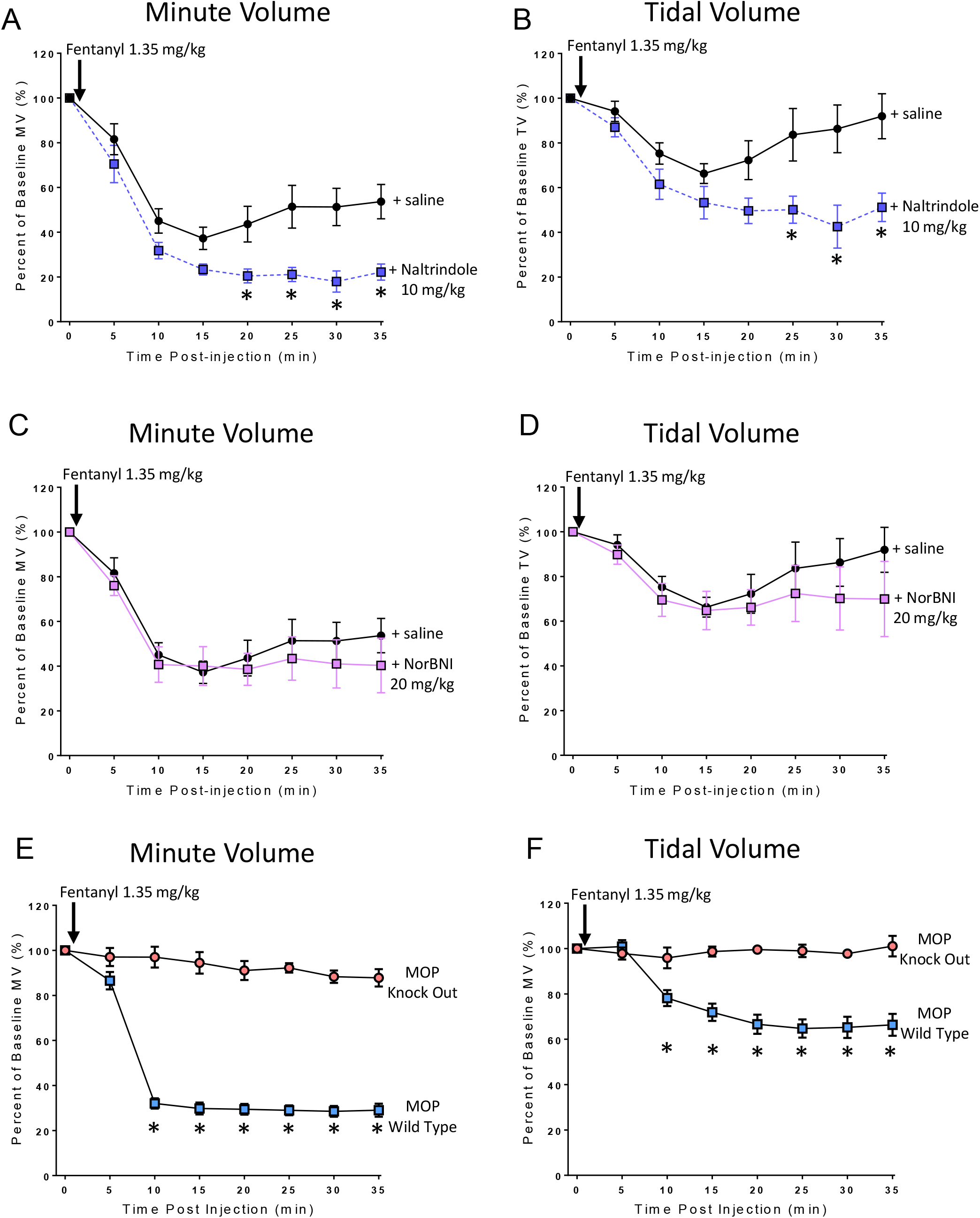
Fentanyl depression of respiration results from activation of μ opioid receptors. **A & B.** Pre-treatment with naltrindole (10 mg.kg^−1^ i.p.) 20 min prior to fentanyl (1.35 mg.kg^−1^ i.p.) injection did not prevent fentanyl depression of minute volume (**A**) or tidal volume (**B**). Instead naltrindole pre-treatment significantly enhanced fentanyl depression of both minute volume and tidal volume. **C & D.** Pre-treatment with NorBNI (20 mg.kg^−1^ i.p.) 24 h prior to administration of fentanyl did not alter fentanyl-induced depression of minute volume (**C**) or tidal volume (**D**). **E.** Administration of fentanyl (1.35 mg.kg^−1^ i.p.) significantly depressed minute volume in wild-type background strain mice, but there was no effect in μ opioid receptor knock-out mice [F = 933 (dfn = 1, dfd = 80)]. **F.** Fentanyl (1.35 mg.kg^−1^ i.p.) also depressed tidal volume in wild type mice but there was no effect in μ opioid receptor knock-out mice [F = 170.9 (dfn = 1, dfd = 80)]. All data presented as mean ± s.e.m. Statistical comparison made by 2-way ANOVA with Bonferroni’s comparison. * indicates p<0.05 compared to saline pre-treated mice. n=6 for each group.

Finally, we examined the ability of fentanyl to depress respiration in μ opioid receptor knock-out mice (C57BL/J background strain). Administration of fentanyl (1.35 mg.kg^−1^ i.p.) to the wild type background strain of mice depressed minute volume by ~80% (Fig. 5E) and depressed tidal volume by ~20% (Fig. 5F). Administration of the same dose of fentanyl to μ opioid receptor knock-out mice produced no depression of either minute or tidal volume (Fig. 5E and F).

### Interactions between heroin, morphine and fentanyl

Opioid users are not thought to use fentanyl as their primary drug of choice, rather they predominantly use heroin to which fentanyl has been added to enhance the “quality” of a given batch of heroin (Ciccarone, 2009; Dasgupta *et al.*, 2013). They are therefore likely to have already developed some degree of tolerance to heroin. We therefore investigated the degree of cross tolerance to fentanyl produced by prolonged pre-treatment with morphine, the main active breakdown product of heroin. In control mice that were implanted with a pump containing saline an acute challenge with morphine (10 or 90 mg.kg^−1^ i.p.) produced respiratory depression of 40% and 60% respectively, whereas in mice that had received prolonged pre-treatment with morphine the response to the 10 mg.kg^−1^ challenge dose of morphine was completely abolished and that to 90 mg.kg^−1^ markedly attenuated (Fig. 6A & B). These data demonstrate that the morphine pre-treatment had produced significant tolerance. In contrast morphine pre-treated mice showed significantly less cross tolerance when challenged with fentanyl (Fig. 6C - F). At the lower challenge dose of fentanyl (0.15 mg.kg^−1^) the depression of respiration was partially reduced but to a lesser extent than the equipotent challenge dose of morphine (Fig. 6C & E). At the higher challenge dose of fentanyl (1.35 mg.kg^−1^) respiratory depression was the same as in non-morphine treated animals (Fig. 6D & F).

**Figure 6.**
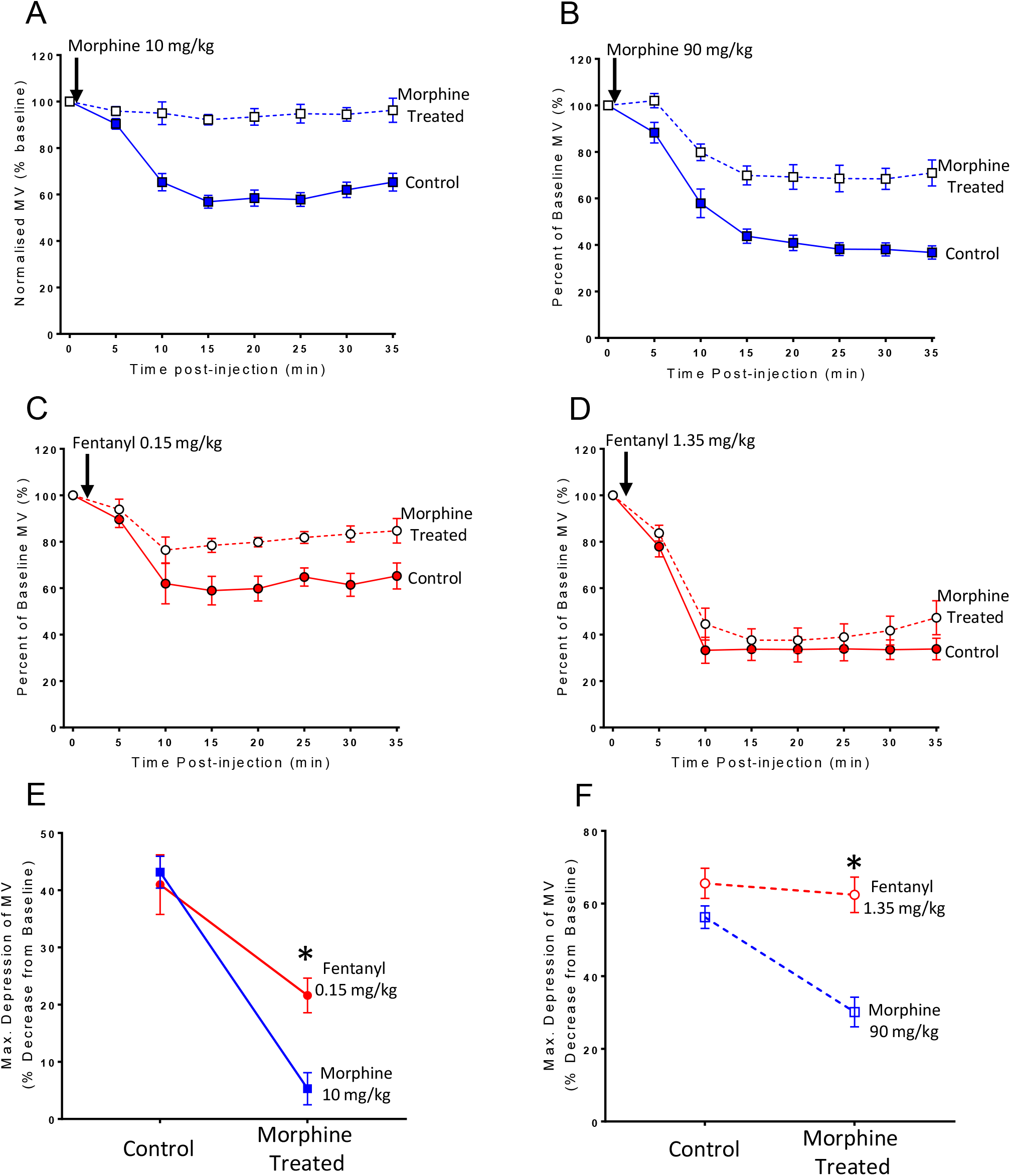
Cross tolerance to fentanyl following prolonged morphine administration. **A-B.** Acute morphine challenge (10 or 90 mg.kg^−1^ i.p.) induced less respiratory depression in morphine-treated mice compared to saline-treated controls. **C.** Acute fentanyl challenge (0.15 mg.kg^−1^ i.p.) induced less respiratory depression in morphine-treated mice compared to saline-treated controls. **D.** Acute fentanyl challenge (1.35 mg.kg^−1^ i.p.) induced the same level of respiratory depression in morphine-treated mice as was observed in saline-treated controls. **E-F.** The depression of respiration by fentanyl challenge (0.15 and 1.35 mg.kg^−1^) was not reduced to the same extent as that by morphine challenge (10 or 90 mg.kg^−1^) in morphine-treated animals. All data presented as mean ± s.e.m. Statistical comparison made by 2-way ANOVA with Bonferroni’s comparison. **E** [F = 3.86 (dfn = 1, dfd = 24)] **F** [F = 25.7 (dfn = 1, dfd = 24)] * indicates p<0.05 compared to morphine. n=7 for each group.

## DISCUSSION

In the present study we observed that in the mouse fentanyl more rapidly depressed respiration than heroin and that with fentanyl the depression of respiration involved both a decrease in the frequency of breathing and a decrease in tidal volume whereas with heroin only at the highest dose was a small effect on tidal volume observed. The decrease in tidal volume produced by fentanyl may be due to an increase in muscle stiffness and this would be supported by fentanyl reducing locomotor activity unlike heroin, which increased locomotor activity. We also observed that the depression of respiration by fentanyl required higher doses of the opioid antagonist naloxone to be reversed than did the depression induced by morphine, the active breakdown product of heroin. In contrast diprenorphine reversed fentanyl and morphine depression of respiration to the same extent. The depression of respiration by fentanyl is mediated by the μ opioid receptor because it was not observed in μ opioid receptor knock-out mice and in wild type mice it was reversed by naloxone (μ, δ and κ opioid receptor antagonist) but was unaffected by naltrindole or NorBNI, selective inhibitors of δ and κ opioid receptors respectively. Finally, prolonged pre-treatment of mice with morphine produced less cross tolerance to fentanyl than the tolerance to morphine itself. From these data we conclude that fentanyl is not simply a potent opioid agonist but that it depresses respiration in ways that heroin and morphine do not. These additional effects may contribute to the high lethality of fentanyl.

The higher *in vivo* potency of fentanyls compared to other opioids is well documented as a factor in their lethality. However, we observed in this study that fentanyl was not only approximately 70x more potent than morphine and heroin when depressing mouse respiration, but it also had a significantly faster rate of onset to depress respiration. A fast rate of onset has been reported in fentanyl-using human populations with fentanyl suggested to produce lethal respiratory depression as quickly as 2 min following injection (Burns *et al.*, 2016; Green *et al.*, 2016). This would make effective intervention with naloxone in fentanyl overdose more difficult to achieve.

We have previously reported that in the mouse the depression of respiration by several μ opioid receptor agonists, morphine, oxycodone and methadone, results from a reduction in respiratory rate and at the doses tested does not involve a decrease in tidal volume (depth of breathing) (Hill *et al.*, 2018; Hill *et al.*, 2016; Withey *et al.*, 2017). In the present study however, we observed that fentanyl depressed both respiratory rate and tidal volume whereas at equi-effective doses, the effect of heroin was primarily due to a decrease in respiratory rate. The decrease in tidal volume induced by fentanyl may be indicative of respiratory muscle stiffness inducing a decrease in muscle compliance that attenuates the expansion of the lungs and therefore effects a decrease in mouse tidal volume. In man, intravenous administration of fentanyl and alfentanil produces skeletal muscle rigidity resulting in stiffness of the chest wall (Benthuysen *et al.*, 1986; Streisand *et al.*, 1993; Waller *et al.*, 1981). Brain micro injection studies in rats have implicated several brain regions - locus coeruleus, basal ganglia, nucleus raphe pontis and periaqueductal grey - as sites of action of fentanyls to induce muscle rigidity (Blasco *et al.*, 1986; Lui *et al.*, 1989; Lui *et al.*, 1990; Slater *et al.*, 1987; Weinger *et al.*, 1991; Widdowson *et al.*, 1986) and have shown that it is mediated by activation of μ, and not δ or κ opioid receptors (Vankova *et al.*, 1996). It is likely therefore that in man intravenous injection of fentanyl results in both a decreased drive to breathe and a mechanical resistance to breathing both of which would contribute to overdose death (Burns *et al.*, 2016). Signs of muscle rigidity have been observed in opioid injectors who have presumably injected illicit fentanyls at a supervised drug injection facility in Vancouver (Kinshella *et al.*, 2018).

Our finding that naloxone less readily reversed respiratory depression by fentanyl compared with morphine confirms reports from studies in man that more naloxone may be required to reverse a fentanyl overdose compared to a heroin overdose (Fairbairn *et al.*, 2017; Lynn *et al.*, 2018; Peterson *et al.*, 2016; Schumann *et al.*, 2008; Somerville *et al.*, 2017). In our study we allowed the depression of respiration by each agonist to reach maximum before naloxone was administered to mimic how the drugs would be administered in an overdose situation. Lower sensitivity of fentanyl to antagonism by naloxone cannot be explained simply by fentanyl having high affinity for the μ opioid receptor given that under competitive conditions the degree of antagonism does not depend on the affinity of the agonist but only upon the affinity and concentration of the antagonist (Rang *et al.*, 2016). Furthermore, the lipophilic opioid antagonist, diprenorphine reversed both fentanyl and morphine depression of respiration equally. Potential explanations include non-equilibrium binding conditions for lipophilic ligands in the intact organism, or perhaps that fentanyl and diprenorphine which are potent, highly lipophilic opioid ligands bind within the ligand binding pocket of the μ opioid receptor in a manner distinct from morphine and naloxone.

We have previously demonstrated that tolerance develops to the respiratory depressant effects of μ opioid receptor agonists such as morphine, oxycodone and methadone (Hill *et al.*, 2018; Hill *et al.*, 2016). In the present study we observed that when morphine pre-treated mice were challenged with either fentanyl or morphine the degree of cross tolerance to fentanyl was less than the tolerance to morphine itself. A similar reduced level of cross tolerance between fentanyl and morphine was observed in a study of opioid-induced locomotor activity (Brase, 1986). Cross tolerance develops between drugs acting at the same receptor, but the degree of cross tolerance will depend upon agonist intrinsic efficacy. Fentanyl has higher intrinsic efficacy (McPherson *et al*., 2010) and so needs to occupy a smaller proportion of the available receptors to produce its response. It will therefore be less affected by the loss of μ opioid receptor function - from either receptor desensitization, internalization or degradation – that underlies tolerance. The implication being that with opioid drug users fentanyl will be able to ‘break through’ tolerance induced by heroin, oxycodone and methadone and produce respiratory depression.

## CONCLUSIONS

Our studies in mice indicate that in overdose a number of factors may contribute to the high lethality of fentanyl. These include its high potency, rapidity of onset of action, depression of rate and depth of respiration, lower sensitivity to reversal by naloxone and reduced cross tolerance to other abused opioids.

## Non standard abbreviations

AUC: Area under the curve
NorBNI: nor-Binaltorphimine

## Author contributions

R.H., W.L.D., E.K. and G.H. participated in research design.

R.H. and R. S. performed the experiments.

R.H., R.S. and G.H. analysed the data.

R.H., W.L.D., E.K. and G.H. participated in writing the manuscript.

## Acknowledgements

We thank Ruby Fletcher for assistance with respiratory data analysis.

